# Haplotype assembly without parental sequencing: Genotype-based trio-binning (GT-Trio)

**DOI:** 10.64898/2026.06.08.729486

**Authors:** Thea Johanna Hettasch, Arne Bjørke Gjuvsland, Matthew Peter Kent, Harald Grove, Dag Inge Våge

## Abstract

Trio-binning is a robust method for haplotype-separated assembly, providing the most accurate representation of diploid genomes including complex and haplotype-specific variation. Conventional *trio-binning* methods depend on parental short-read sequences to differentiate offspring reads originating from the maternal and paternal haplotypes. Here, we present a genotype-based trio-binning pipeline (GT-Trio) which reconstructs parent sequences based on phased parental genotypes and uses this as an alternative source of parental information for haplotype assembly. The GT-Trio pipeline was applied to assemble the maternal and paternal haplotypes of three Norwegian Red cattle individuals, using either imputed sequence variants or SNP array genotypes as input. Haplotypes assembled with GT-Trio using sequence variants as parental input demonstrated high phasing accuracy, similar to that achieved with conventional trio-binning. Using lower-density panels of array SNPs as input resulted in a decrease in phasing accuracy, but still substantially improved haplotype resolution compared to trio-free assembly. We show that the use of high-density parental information from short-reads or sequence variants increases the level of fragmentation in repeat-rich and low heterozygosity regions. Using lower-density parental information from array resolves the contiguity in these regions, but at the expense of phasing accuracy. Overall, GT-Trio provides a robust and scalable framework for haplotype assembly without parental sequencing and is available at https://github.com/theahettasch/GT-Trio.

## Background

Haplotype-separated genome assemblies provide the most accurate representation of diploid genomes. They preserve all complex and haplotype-specific regions, which can easily be misrepresented in collapsed assemblies (1–4). The trio-binning method, introduced by Koren et al., uses sequencing data from parents to separate and assemble the maternal and paternal haplotypes of an offspring individual (5). In trio-binning, haplotype-specific k-mers (hapmers) are identified from parental sequences and used to separate and assemble offspring reads originating from each individual haplotype. The trio-binning method was originally implemented as a pre-binning strategy, where offspring reads were separated into paternal and maternal bins prior to assembly (5). However, this pre-binning strategy can be prone to mispartitioning of reads in highly complex areas. Hence, a more robust graph-based assembly strategy (implemented in Hifiasm) was later developed to overcome this issue (6). Graph-based trio-binning omits the pre-binning step by building a string graph based on local overlap between offspring reads and using hapmer labelling to infer maternal and paternal haplotype paths from this graph (6).

Existing implementations of trio-binning require short-read sequencing data from parental genomes, representing a possible bottleneck for haplotype assembly, as it increases costs, requires additional laboratory work and depends on the availability of parent individuals for sampling. Although alternative approaches integrating Strand-Seq (7) or Hi-C (8) can generate haplotype-resolved assemblies without parental data, these methods also require the generation of additional datatypes. Across livestock and some aquacultural species, large scale SNP genotyping and imputation is routinely performed as a part of genomic selection programs, providing phased genotype data at population scale (9–13). In populations where such resources are established, these datasets provide a readily available source of parental genotype information that could be leveraged for trio-binning and enable haplotype assembly for a larger number of individuals without additional sequencing of parent genomes.

In this study we present a genotype-based trio-binning pipeline called GT-Trio, which uses phased genotypes, instead of short-read sequences, from parents to assemble offspring haplotypes with graph-based trio-binning. Using data from three Norwegian Red (NR) cattle trios, we apply the GT-Trio pipeline to assemble offspring haplotypes using both imputed sequence variants and SNP array genotypes as parental input. We evaluate how the GT-Trio pipeline impacts assembly quality, contiguity, and accuracy of haplotype separation compared to conventional trio-binning.

## Methods

### Genotype-based trio-binning pipeline (GT-Trio)

The genotype-based trio-binning pipeline (GT-Trio) was developed to automate haplotype assembly using parental genotypes as input for trio-binning. The pipeline is implemented as a Snakemake workflow and is freely available at https://github.com/theahettasch/GT-Trio.

GT-Trio takes (i) a high-quality reference genome (FASTA), (ii) phased parental genotypes (VCF), and (iii) offspring ONT long-reads (FASTQ) as input and outputs two chromosome-level assemblies representing the maternal and paternal offspring haplotypes. The pipeline can take any dataset of phased parental genotypes, including SNPs, INDELs and larger SVs. The pipeline executes the following five steps implementing different software tools (Figure 1):

1. Parental sequence reconstruction: Parent-specific sequences are reconstructed by modifying a high-quality reference genome according to a set of phased parental genotypes with BCFtools consensus (14).
2. Parental read simulation: Paired-end 150 bp reads are simulated from the parent-specific sequences with SAMtools wgsim (14).
3. Parental k-mer dictionary construction: Parental k-mer dictionaries are created from simulated parental reads with Yak (15).
4. Haplotype-resolved assembly: ONT offspring long-reads are assembled into contig-level haplotypes with Hifiasm (6, 16) using the parental k-mer dictionaries as input for trio-binning.
5. Reference-guided scaffolding: Contigs assembled with Hifiasm are combined into chromosome-level scaffolds with RagTag (17).

**Figure 1:**
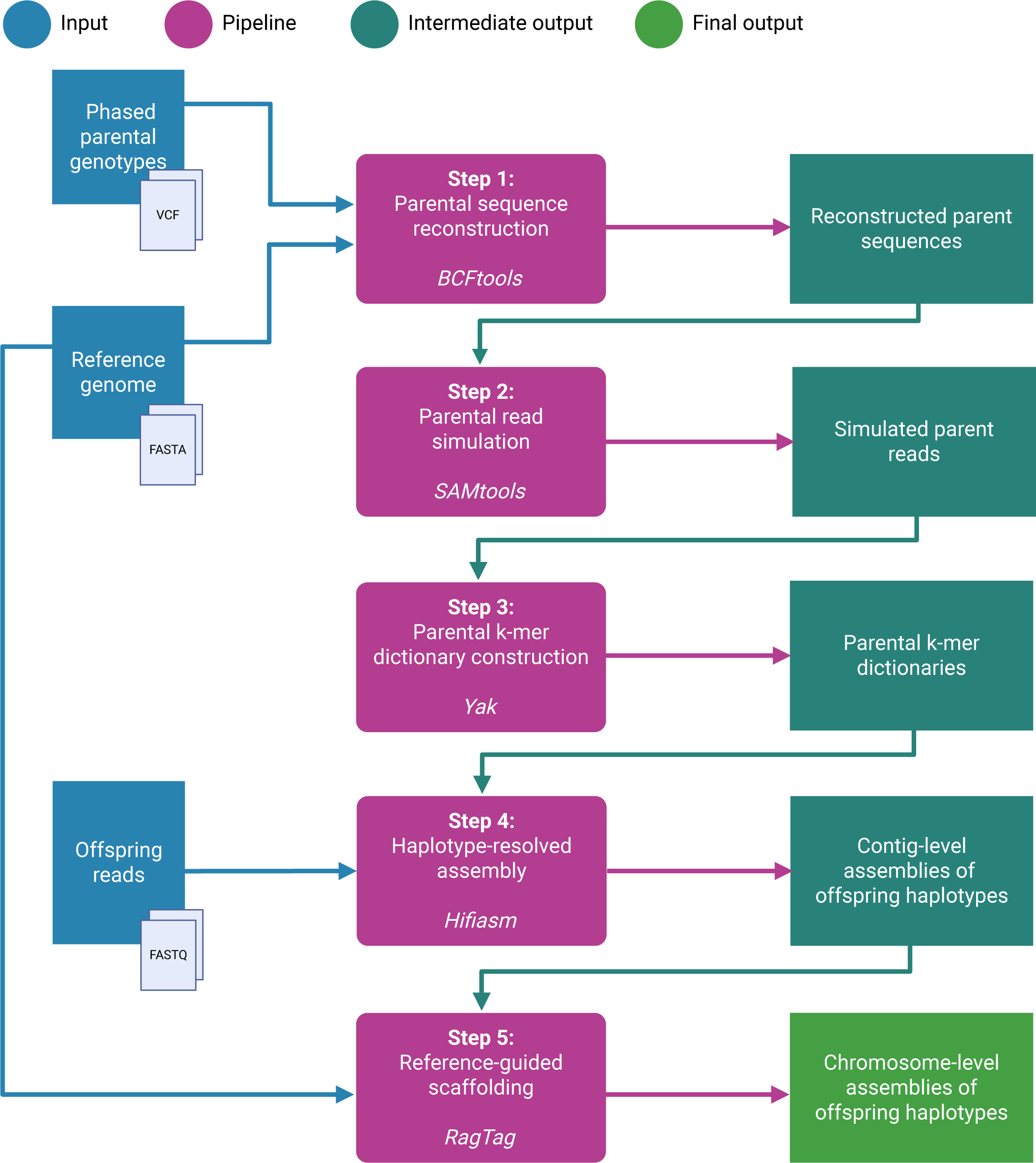
GT-Trio pipeline workflow. Flowchart illustrating the GT-Trio pipeline workflow. GT-Trio performs five steps implementing different software tools. The pipeline requires three input files: a reference genome (FASTA), phased parental genotypes (VCF) and a set of offspring ONT reads (FASTQ). Intermediate output files are produced in each pipeline step and used as input in the following step. The final output of the pipeline is two chromosome-level assemblies representing offspring haplotypes.

Detailed descriptions of pipeline steps, required input files and parameters are provided in the pipeline documentation: GT-Trio/README.md.

### Validation of GT-Trio pipeline in Norwegian Red cattle

The GT-Trio pipeline was applied to construct haplotype-resolved assemblies using data from three Norwegian Red (NR) cattle trios. Nanopore (ONT) long reads (53-58X) from three NR trio offspring individuals, described in a previous study (18), were used in this study and are available on the European Nucleotide Archive (ENA) (Table 1). Reads shorter than 1000 bp were removed with Filtlong (19).

**Table 1:** ONT long-read datasets.

| Trio | Individual | Sequencing platform | ENA Accession | Coverage | Read N50 | Read # |
| --- | --- | --- | --- | --- | --- | --- |
| 1 | Offspring | ONT PromethION | ERX15390475 | 53X | 32 kb | 7.5 M |
| 2 | Offspring | ONT PromethION | ERX15390514 | 56X | 32 kb | 7.8 M |
| 3 | Offspring | ONT PromethION | ERX15390528 | 58X | 31 kb | 8.6 M |

ONT long-read sequencing datasets from NR trio offspring individuals used for assembly in this study. All read sets are
available on ENA under the accession number.

Phased genotypes for NR trio parents were imputed from array to sequence level. Imputation from 50K to 777K density was performed routinely by the NR breeding company Geno as described in Nordbø et al. (20). Subsequently, genotypes for ∼255K NR individuals, including all trio parents, were imputed from 777K to sequence level using a reference population of 633 short-read sequenced NR individuals. The imputation reference was created by calling SNPs and INDELs from Illumina short reads using HaplotypeCaller from the GATK software package (v4.5) (21). Phasing and imputation was done with Beagle v.5.4 (22). Variant positions were based on a near-complete NR reference (GCA_963921495.1). All autosomes were included.

The maternal and paternal haplotypes of the NR offspring were assembled with GT-Trio. The NR reference genome (GCA_963921495.1) was downloaded from ENA and given as input together with the imputed parental genotypes and the filtered offspring ONT reads. Applied pipeline parameter settings are provided in Methods S1 (File S1). To explore the effect of parental genotype density on assembly quality, the following subsets were extracted from the imputed dataset and used respectively as input to GT-Trio:

- All imputed sequence variants
- SNPs on the 777K array (Illumina BovineHD)
- SNPs on the NR-specific 50K array (Illumina NRF v2)
- SNPs on the multibreed bovine 50K array (Illumina BovineSNP50 v2.0)

To explore the sensitivity of GT-Trio to the use of a reference genome from a more distantly related cattle breed, the Hereford reference (ARS-UCD1.2, GCA_002263795.2) was used to assemble NR haplotypes with a panel of 777K SNPs with Hereford-specific genomic coordinates.

### Conventional trio-binning and trio-free assembly of NR offspring haplotypes

To benchmark the GT-Trio pipeline, NR offspring haplotypes were also assembled with conventional trio-binning (Hifiasm), using parental Illumina short-reads as input, and trio-free assembly using no parental input (Methods S2, File S1). Illumina short-reads (28-42 X) from NR trio parents, are available on ENA (Table 2).

**Table 2:** Illumina short-reads datasets.

| Trio | Individual | Sequencing platform | ENA Accession | Coverage |
| --- | --- | --- | --- | --- |
| 1 | Sire | Illumina NovaSeq 6000 | ERX15390479, ERX15390480 | 29X |
| 1 | Dam | Illumina NovaSeq 6000 | ERX15390481 | 33X |
| 2 | Sire | Illumina NovaSeq 6000 | ERX15390517, ERX15390518 | 28X |
| 2 | Dam | Illumina NovaSeq 6000 | ERX15390520 | 39X |
| 3 | Sire | Illumina NovaSeq 6000 | ERX15390530 | 28X |
| 3 | Dam | Illumina NovaSeq 6000 | ERX15390532 | 42X |

Illumina short-reads sequencing datasets from NR parents used as input for conventional trio-binning. All read sets are available on ENA under the accession number.

### Evaluation of assembly quality

Haplotype assemblies were evaluated in terms of size, contiguity, completeness and accuracy of haplotype separation. All autosomes were considered for evaluation of chromosome-level assemblies.

The assembly size and contig N50 values were calculated with gfastats (23). As we move towards near-gapless assemblies, the N50 value loses some of its significance as a measure of contiguity as it only considers half of the assembly in its calculation. So, to complement this, we also evaluated the contig/chromosome (CC) ratio (24). Compleasm was used to assess the presence of universal single-copy orthologs (BUSCO genes) against the Mammalia_od10 database (9226 genes) (25).

Accuracy of haplotype separation was evaluated based on the presence of haplotype-specific k-mers (hapmers) in the final assemblies. K-mer dictionaries (k = 21) were generated from offspring ONT reads and parental Illumina short-reads with Meryl (26) and hapmers were identified as the subset of offspring k-mers (KO) present exclusively in either the paternal (KP) or maternal (KM) set of k-mers:

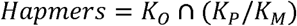

These hapmers were used as a validation set to assess the accuracy of haplotype separation in all assemblies constructed with GT-Trio, conventional trio-binning and trio-free methods. Paternal and maternal hapmers were mapped to the paternal and maternal haplotype assemblies with Merqury (26). Ideally, any maternal hapmers should only be present in the maternal haplotype assembly, while any paternal hapmers should only be present in the paternal haplotype assembly. Deviations from this were quantified using the Hamming error rate:

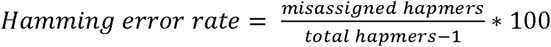

See Methods S3 (File S1) for details on how Meryl and Merqury were run to assess accuracy of haplotype separation.

### Visualization of synteny between haplotypes

Synteny between pairs of paternal and maternal haplotypes was identified with Syri (27) based on assembly alignments generated with minimap2 (28). Syntenic regions and rearrangements were visualised with plotsr (29).

### Repeat annotation of assembly gaps and phasing errors

To explore associations between repetitive motifs and assembly errors, we analysed the repeat content of genomic regions adjacent to observed contig breaks and phasing errors. To enable comparison of errors across assemblies, contig breakpoints and hapmer error positions were projected onto the NR reference genome using the following approaches. Contig breakpoints were retrieved directly from the RagTag scaffolding output, which reports contig alignments relative to the NR reference. In contrast, hapmer error positions were lifted over from individual haplotype assemblies to the NR reference using CrossMap (30). Lift-over was performed with CHAIN files generated from whole-genome alignments between each haplotype assembly and the NR reference, with minimap2 (28) (Methods S4, File S1).

A *de novo* repeat annotation was generated for the NR reference with RepeatModeler (31) and RepeatMasker (32). The NR reference genome was followingly soft masked with BEDtools (33) and given as input to Biser (34) for annotation of Segmental Duplications (SD) (Methods S4, File S1). The acrocentric centromeres of reference autosomes were annotated using the bovine satellite enrichment approach described by Hettasch et al. (18). Estimated centromere lengths are provided in Figure S1 (File S2). Contig break and hapmer error positions were intersected with the reference repeat annotation using a custom R script (Methods S4, File S1). Positions within ± 100 bp of an annotated repeat were considered repeat-associated.

## Results

### Haplotype-resolved assembly of Norwegian Red cattle genomes with GT-Trio

The GT-Trio pipeline was applied to construct haplotype assemblies using data from three Norwegian Red (NR) cattle trios. Phased parental genotypes were imputed from array to sequence for all trio parents, achieving 20 946 844 phased and genotyped sequence variants. Across the panel of imputed sequence variants, the median dosage-R2 (DR²) was 0.99, with 76.7% having DR² > 0.8. Imputation quality was lower for very rare variants (AF < 0.005; mean DR² = 0.32) but remained high for more frequent variants (mean DR² = 0.95 for AF ≥ 0.01 and 0.97 for AF ≥ 0.05). These results indicate that the imputed genotypes used as input to GT-Trio were generally of high quality. The full set of sequence variants included 626 080 SNPs from the 777K (Illumina BovineHD) array, 48 659 SNPs from the 50K NR-specific (Illumina NRF v2) array and 44 813 SNPs from the 50K multibreed bovine (Illumina BovineSNP50 v2.0) array. The full set of sequence variants as well as subsets of SNP array genotypes were used as parental input for GT-Trio. For comparison, haplotypes were also assembled using conventional trio-binning, with parental Illumina short-reads as input and trio-free assembly without parental input.

### Density of phased parental genotypes impacts hapmer recovery

Haplotype-specific k-mers (hapmers) represent unique paternal and maternal sub sequences and are used in trio-binning to label offspring reads originating from the maternal and paternal haplotypes (Figure 2a). In a panel of parental genotypes, only variants diverging between parents can be informative for hapmer identification (Table S1 in File S3). The panel of sequence variants covered the highest number of informative SNPs. However, the relative proportion of informative SNPs is higher in array-based panels, as they are designed to capture genetic variation (Table S2 in File S3). The number of hapmers recovered from sequence variants is similar to the number of hapmers recovered from parental Illumina short-reads (Figure 2b; Data S1 in File S4), suggesting that the panel captures most of the heterozygosity between parental haplotypes. In contrast, parental sequences reconstructed from array SNPs show a substantial reduction in the number of hapmers that can be recovered as only a subset of variants is included on the array (Figure 2b; Data S1 in File S4). Sequences reconstructed from SNPs on the NR-specific 50K array recover more hapmers than sequences reconstructed from SNPs on the multibreed bovine 50K array (Figure 2b; Data S1 in File S4), confirming that the breed-customised array covers more informative variation.

**Figure 2:**
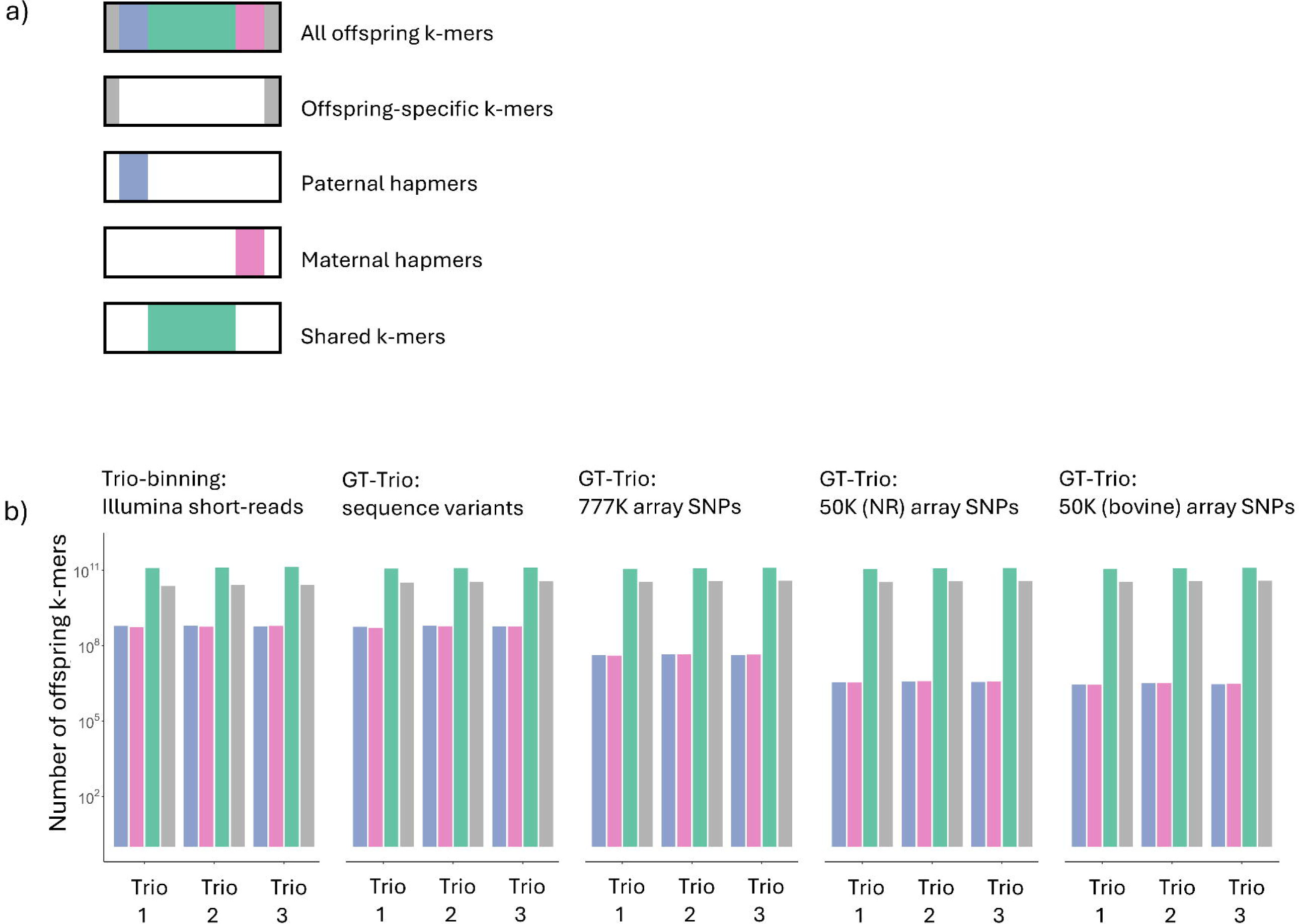
Categorisation of offspring k-mers a) Categorisation of k-mers found in offspring reads. Offspring-specific k-mers (grey) are uniquely found in the offspring read set. Haplotype-specific k-mers (hapmers) represent offspring k-mers that are uniquely found in either the paternal (blue) or maternal (pink) k-mers. Shared k-mers (green) represent the overlap of offspring, paternal, and maternal k-mers. b) The number of offspring-specific k-mers, shared k-mers and paternal and maternal hapmers found in NR trio offspring reads. Offspring k-mers are categorised based on their overlap with parental k-mers derived from parental Illumina short-reads or parent sequences reconstructed with GT-Trio using all imputed sequence variants, as well as subset of 777K bovine array SNPs, 50K NR-specific array SNPs and 50K multibreed bovine array SNPs as input.

### Assembly quality

Haplotypes assembled with GT-Trio using parental sequence variants achieved a contig- and chromosome-level assembly size similar to haplotypes assembled with conventional trio-binning, while the use of lower-density array SNPs resulted in an increase in assembly size (Figure 3a,b; Data S2, S3 in File S4). In haplotypes assembled with GT-Trio, a pronounced bias in assembly size is observed between haplotypes (Figure 3a; Figure S2 in File S2). The surplus of sequence observed in the paternal haplotypes is mainly contained in highly repetitive contigs, which end up as unplaced contigs after scaffolding against the NR reference (Figure S4 in File S2), suggesting that these sequences represent highly repetitive sequence missing from the reference (e.g. centromeres and telomeres) or erroneous assembly expansions. Reference-guided scaffolding restores the size balance between haplotypes, and no size bias is observed between the chromosome-level assemblies (Figure 3b; Figure S3 in File S2).

**Figure 3:**
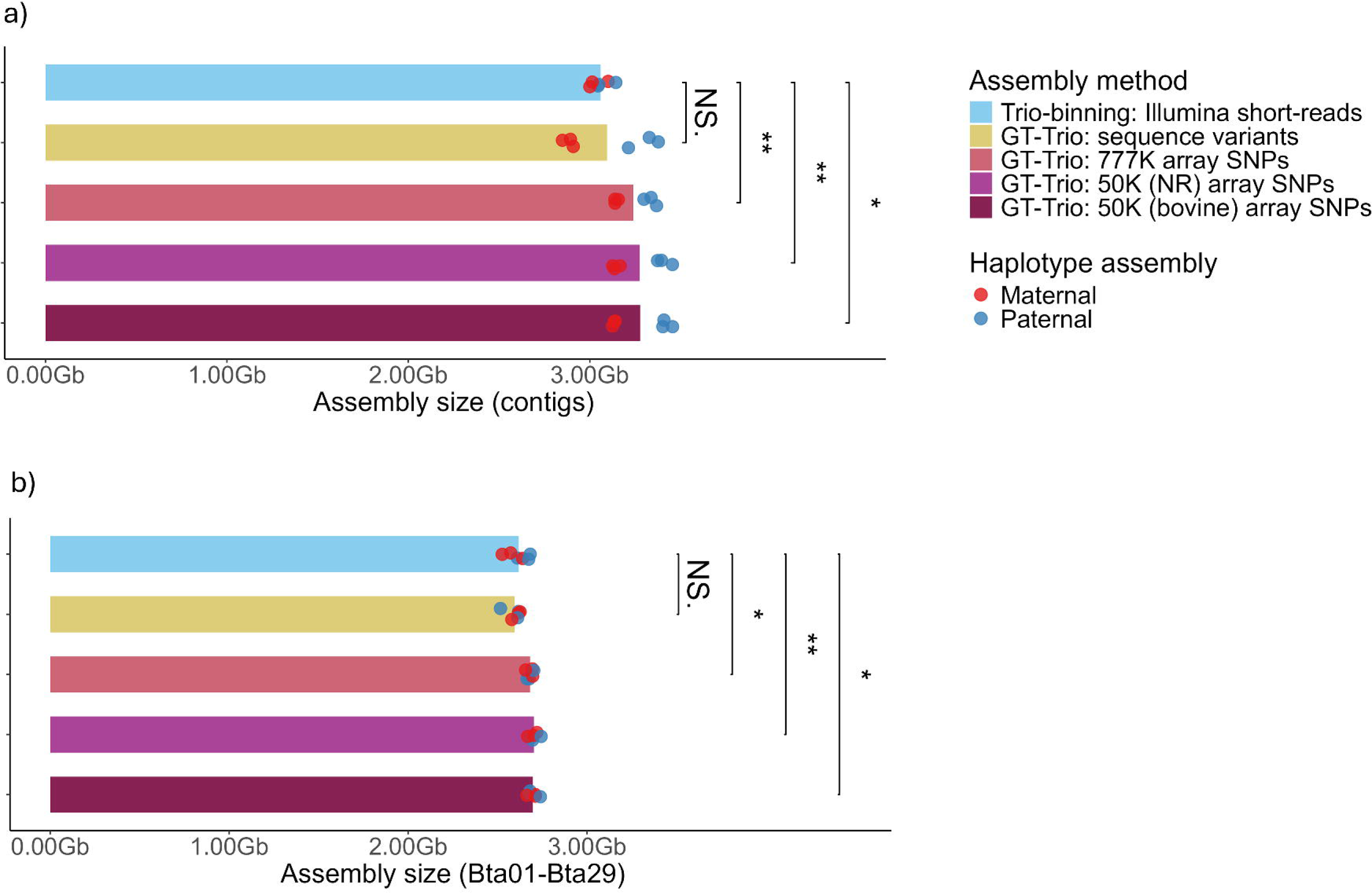
Assembly size Assembly size of haplotypes assembled with conventional trio-binning or GT-Trio. Statistical test: paired t-test, p > 0.05. a) Contig-level assembly size. b) Chromosome-level (Bta01-Bta29) assembly size.

The contig N50 value and CC ratio was used to assess assembly contiguity. No statistically significant differences were observed between haplotypes assembled with conventional trio-binning and GT-Trio (Figure 4a; Data S2, S3 in File S4). However, the mean values indicate that haplotypes assembled with GT-Trio using array SNPs as parental input have higher N50 values and lower CC ratios, suggesting that these assemblies are less fragmented with larger and more contiguous contigs.

**Figure 4:**
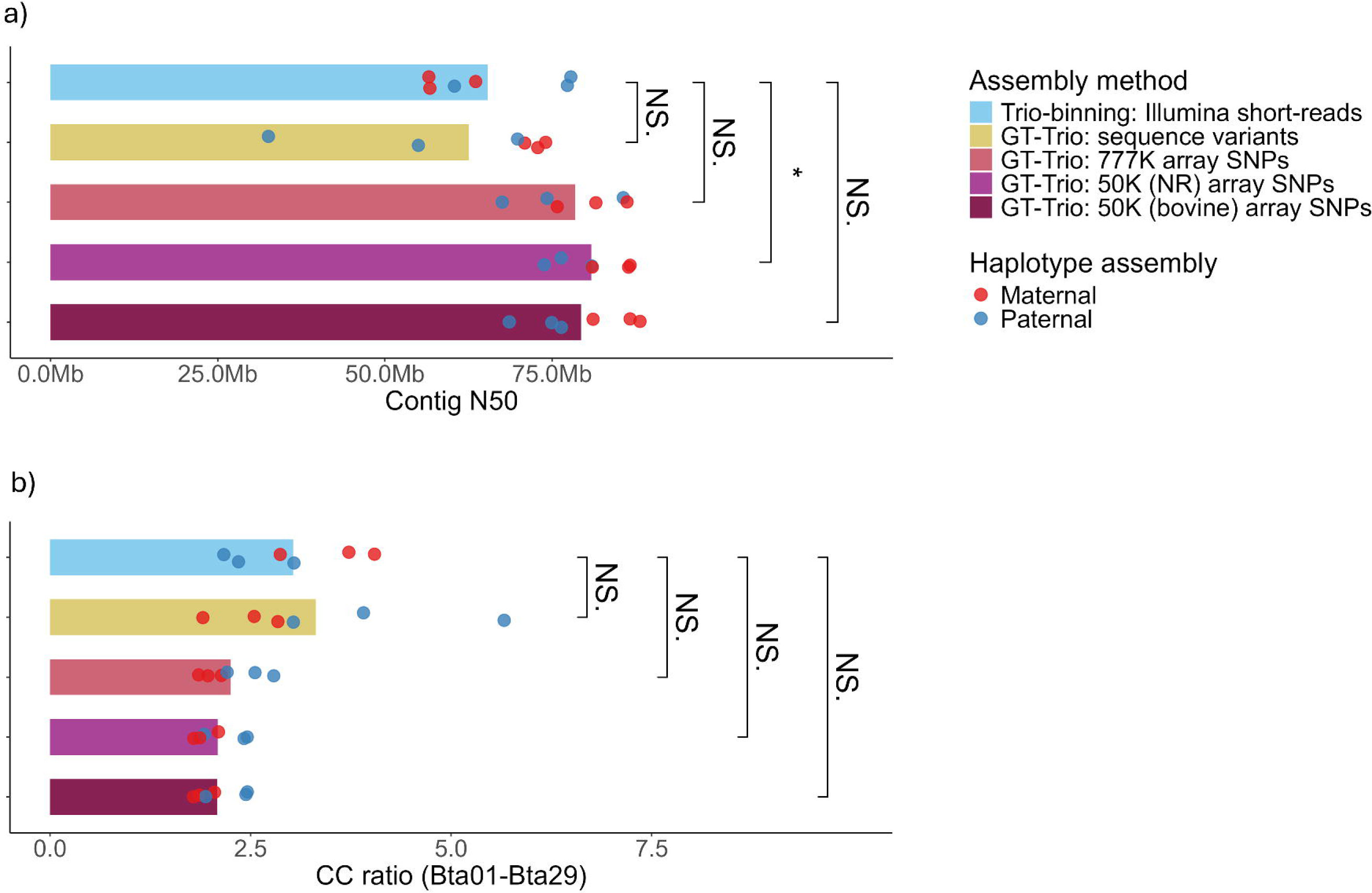
Contig N50 and CC ratios. The contig N50 and CC ratios of NR haplotypes assembled with conventional trio-binning or GT-Trio. Statistical test: paired t-test, p > 0.05. a) Contig N50. b) CC ratio of chromosome-level assemblies (Bta01-Bta29).

BUSCO scores were similar between haplotypes assembled with conventional trio-binning and GT-Trio using sequence variants as parental input (Figure 5a; Data S3 in File S4). Haplotypes assembled with GT-Trio using array SNPs as parental input showed significantly higher BUSCO scores, primarily due to a reduction in fragmented (F) and missing (M) BUSCO genes (Figure 5b,c,d; Data S2, S3 in File S4). In haplotypes assembled with high-density parental information from short-reads or sequence variants there is a markedly high missingness of BUSCO genes.

**Figure 5:**
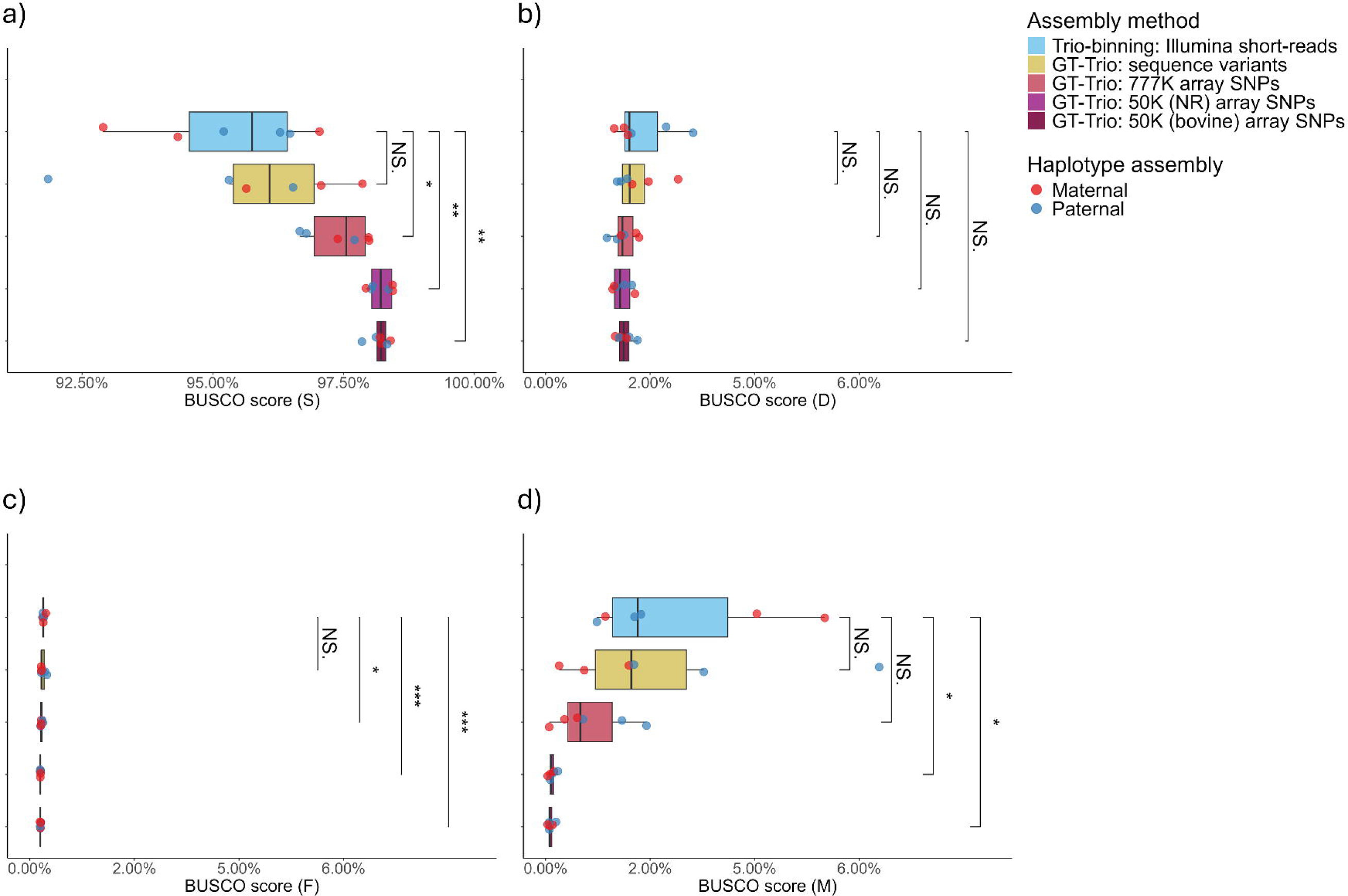
BUSCO scores BUSCO scores of contig-level assemblies. Statistical test: paired t-test, p > 0.05. a) S = Single copy complete genes. b) D = Duplicated complete genes. c) F Fragmented genes – only proportion of gene aligns. d) M = Missing genes.

Synteny alignment of haplotype-pairs reveals larger regions (<Mb) missing from one haplotype and present in the other, which contain the missing BUSCO genes (Figure S5a in File S2). Mapping of hapmers reveal low hapmer density and ambiguous hapmer assignment in these regions, suggesting that they are largely homozygous and should be included in both haplotypes (Figure S5b in File S2). Misassembly in homozygous regions and ambiguous hapmer mapping is generally not observed in haplotypes assembled with GT-Trio using lower-density panels of array SNPs (Figure S5b in File S2).

### Accuracy of haplotype separation

In haplotype assembled with both conventional trio-binning and GT-Trio, the majority of paternal hapmers map to the paternal haplotypes, while the majority of maternal hapmers map to the maternal haplotypes, suggesting good separation of haplotypes (Figure 6a). In contrast, no haplotype separation was achieved with trio-free assembly without parental input, resulting in a mix of both maternal and paternal hapmers mapping to the assemblies (Figure 6a). Accuracy of haplotype, measured by Hamming error rate, did not differ significantly between haplotypes assembled with conventional trio-binning and GT-Trio using sequence variants as parental input (Figure 6b; Data S3 in File S4). Using subsets of array SNPs as input for GT-Trio resulted in a small, but significant increase in Hamming error rate (Figure 6b; Data S3 in File S4). However, these elevated values fall far below the Hamming error rate of haplotypes generated with trio-free assembly, demonstrating how the GT-Trio pipeline achieves significantly better separation of haplotypes with a Hamming error consistently below 5%. Using genotypes from the NR-specific 50K array for assembly with GT-Trio achieves lower Hamming error rates than using the multibreed bovine 50K array SNPs. This is in line with previous observations showing that the NR-specific array covers more informative SNPs than the multibreed bovine array, and followingly recovers more hapmers diverging between parental haplotypes.

**Figure 6:**
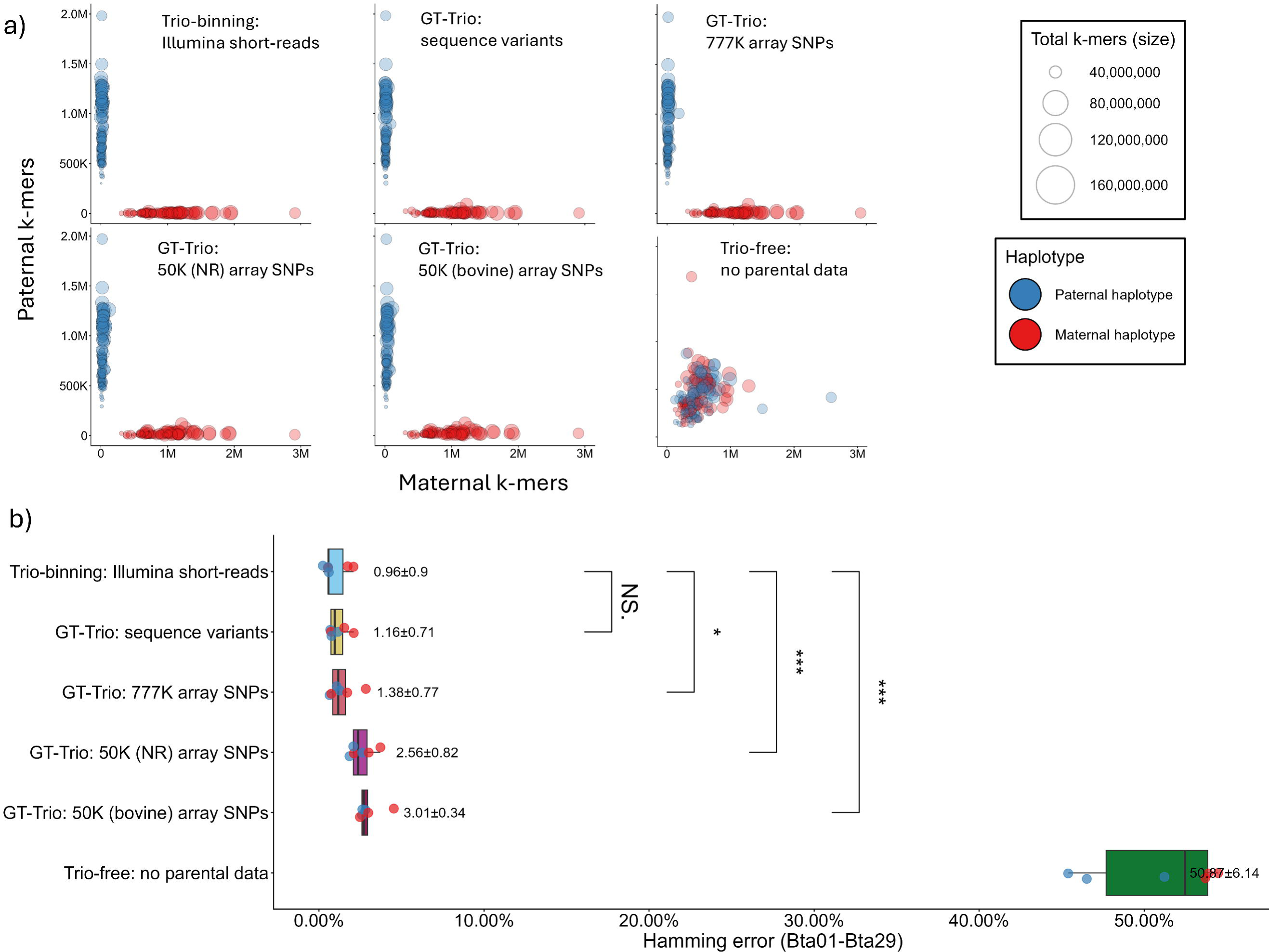
Accuracy of haplotype separation. Accuracy of haplotype separation in assemblies constructed with conventional trio-binning, GT-Trio and trio-free assembly. Statistical test: paired t-test, p > 0.05. a) Maternal and paternal hapmers mapping to contigs in maternal and paternal haplotype assemblies. b) Hamming error rate of maternal and paternal assemblies (Bta01-Bta29).

### Sensitivity to choice of reference genome

The GT-Trio pipeline is dependent on a reference genome as input for reconstruction of parental sequences. To assess the sensitivity of GT-Trio to the choice of reference genome, we assembled NR haplotypes using the more distantly related Hereford reference (ARS-UCD1.2) and a panel of 777K array SNPs with Hereford reference-specific genomic coordinates. Assembly statistics are broadly similar between reference with a modest increase in assembly size and contig N50s with the use of the NR reference, reflecting the larger size and breed-specificity over ARS-UCD1.2 (Data S2 in File S4). There is a slight reduction in accuracy of haplotype separation when using the Hereford reference, with Hamming error rates differing by 0.22% at most (Data S2 in File S4).

### Repeat motifs associated with contig breaks and phasing errors

Haplotypes assembled with conventional trio-binning and GT-Trio showed differences in assembly contiguity and accuracy of haplotype separation. The use of high-density parental information from parental short-reads or sequence variants ensured highly accurate separation of haplotypes but was accompanied by a high level of fragmentation. Using lower density panels of array SNPs increased the assembly contiguity at the cost of less accurate phasing. Results indicate a trade-off between assembly contiguity and haplotype separation which is dependent on the density of parent-specific information provided for trio-binning. The genomic location of the observed contig breaks and hapmer errors, as well as the repeat content of sequences adjacent to these misassemblies were investigated to assess if differences in contiguity and phasing could be attributed to certain hard-to-assemble regions of the genome. There is a higher density of contig breaks in haplotypes assembled with conventional trio-binning and GT-Trio using sequence variants as parental input, compared to haplotypes assembled with GT-Trio using lower density panels of array genotypes as input (Figure 7a). The highest proportion of contig breaks is observed in satellite enriched centromeres and segmental duplications (Figure 7b), across all assembly approaches, which are known to be complex and hard-to assemble regions (35). The increase in assembly fragmentation associated with the use of higher density parental information from short-reads or sequence variants is most prominent in regions with Long Interspersed Nuclear Elements (LINEs), certain unknown repeat types and non-repeated regions (Figure 7b).

**Figure 7:**
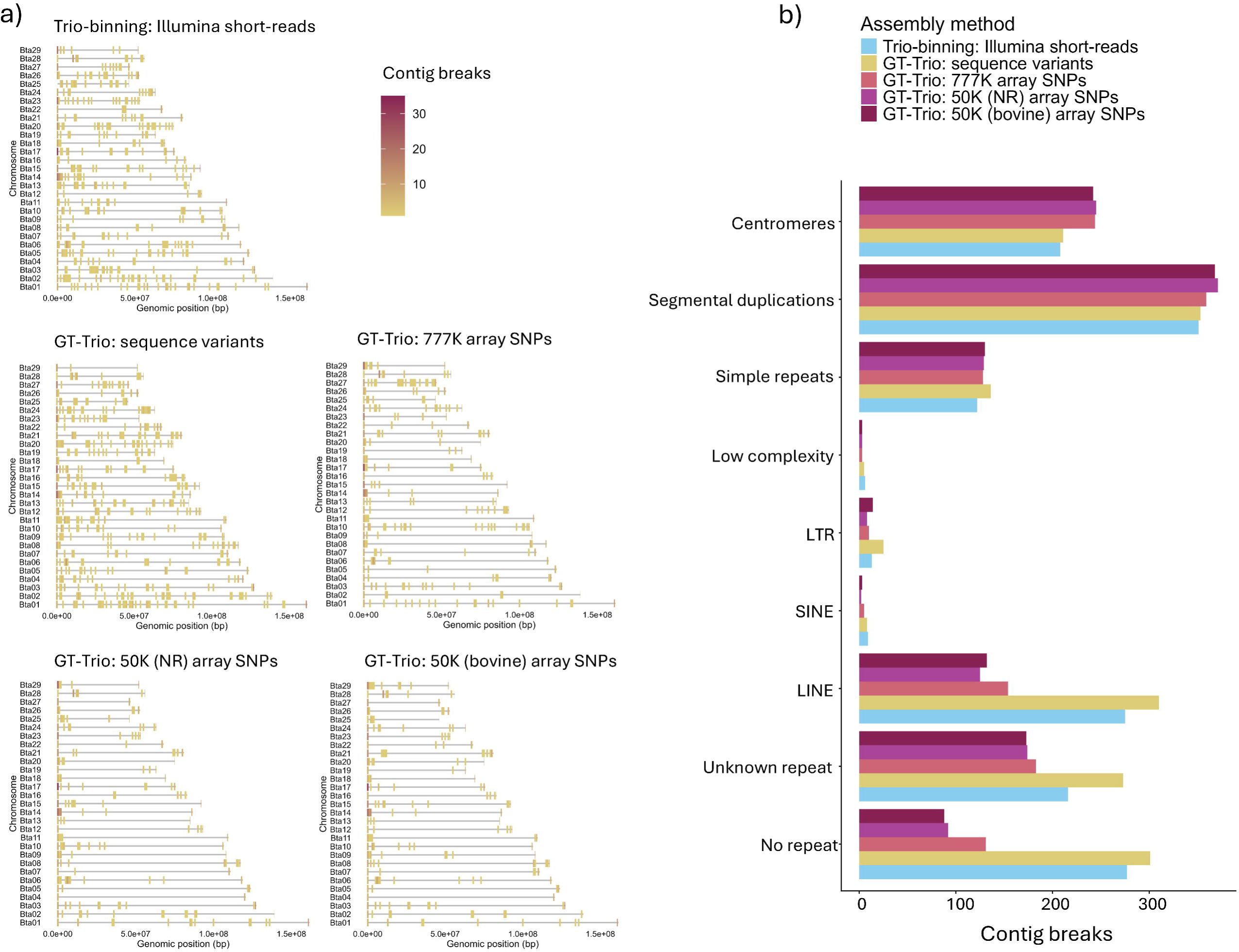
Location and repeat annotation of assembly gaps a) Number of contig breaks within 1 Mb windows across haplotypes assembled with conventional trio-binning and GT-Trio. b) Sequence properties of genomic regions adjacent to contig breaks (± 100 bp).

A higher number of hapmer errors, representing erroneously phased regions, were observed in haplotypes assembled with GT-Trio using array SNPs as parental input (Figure 8a). Using higher density parental information from sequence variants reduced the occurrence of hapmer errors across assemblies. The lowest number of hapmer errors was observed in haplotypes assembled with conventional trio-binning (Figure 8a). Again, difference in hapmer errors were most strongly associated with LINEs, certain unknown repeat regions and non-repeated sequences (Figure 8b), similar to what we observed in our analysis of contig breaks.

**Figure 8:**
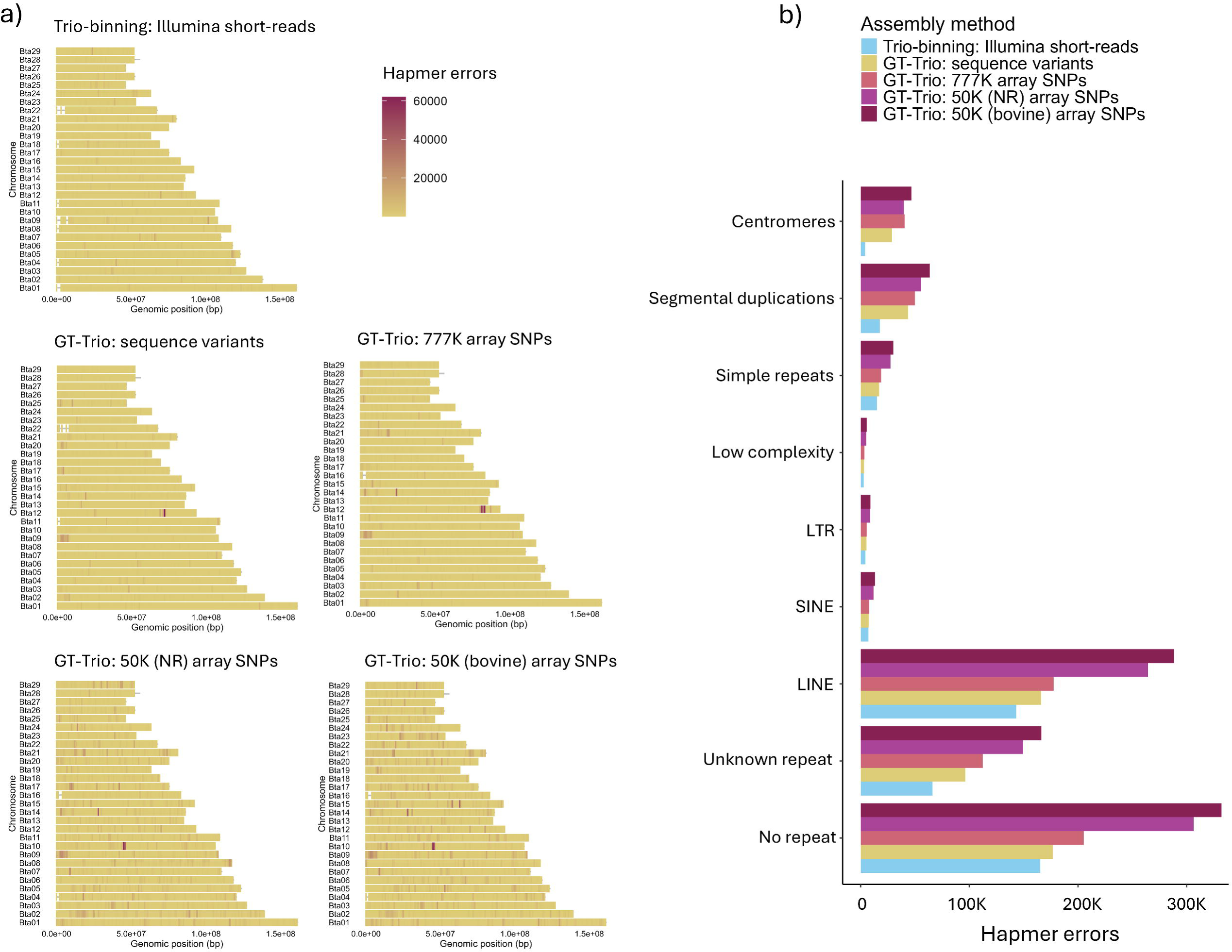
Location and repeat annotation of hapmer errors a) Number of hapmer errors within 1 Mb windows across haplotypes assembled with conventional trio-binning or GT-Trio. Genomic coordinates are based on the NR reference genome. b) Sequence properties of genomic regions adjacent to hapmer errors (± 100 bp).

## Discussion

GT-Trio is a pipeline developed for de novo assembly of haplotypes using phased genotypes as parental input for trio-binning. The pipeline performs parental sequence reconstruction, parental read simulation, k-mer dictionary construction, haplotype assembly and reference-guided scaffolding applying different genomic software tools. GT-Trio utilises parent-specific information contained in phased parental genotypes to separate and assemble the paternal and maternal offspring haplotypes, obviating the need for additional short-read sequencing of parental genomes.

In this study we demonstrate and test the GT-Trio pipeline using data from three Norwegian Red (NR) cattle trios. Phased genotypes were imputed from array to sequence for all trio parents and provided as input to GT-Trio. The accuracy of haplotype-separation achieved with GT-Trio is similar to what we achieve with conventional trio-binning, supporting the use of GT-Trio as an alternative when sequencing of parental genomes is infeasible. Despite the high phasing accuracy, we do observe that NR haplotypes assembled with high-density parental information from short-reads or sequence variants, are compromised in certain regions where larger chunks of sequence are erroneously included in only one of two haplotypes. Results show that these misassemblies occur in regions with ambiguous and low hapmer density, supporting previously reported findings that haplotype assembly can be problematic in highly homozygous regions with weak and ambiguous haplotype-specific signals (36). This complicates successful haplotype assembly of genomes from populations with higher levels of inbreeding, such as the NR trios used in this study. Haplotypes assembled with high density parental information do also show a high level of fragmentation in complex regions including repeats such as LINEs. We suspect that these repeats cause highly complex topological structures in the initial string graph generated by the assembly algorithm, which are further complicated by conflicting or ambiguous haplotype signals. Hifiasm applies a rather conservative trio-binning strategy prioritizing phasing over contiguity in these hard-to-assemble regions. If haplotype signals are conflicting, Hifiasm can remove nodes from the graph, which causes a higher level of assembly fragmentation in these regions (6). Lower-density subsets of variants included on 777K and 50K bovine arrays were also provided as parental input for haplotype assembly with GT-Trio, as they are routinely used for genotyping at a population-scale across many cattle populations. Less parent-specific information is contained in an array-based panels of genotypes compared to the full set of sequence variants, providing less haplotype-specific information for separation of haplotypes during assembly. Followingly, we observed a reduction in the accuracy of haplotype separation in these assemblies. The reduction in phasing accuracy was most prominent in complex genomic regions including repeats and LINEs, which can be more challenging to phase correctly due to complex assembly graph topology. Low representation and genotyping quality of array SNPs in repeated regions can also be contributing to the reduction in accuracy of phasing in these regions. Despite this, the 777K and 50K array SNP panels still provide enough haplotype information to significantly improve phasing over trio-free assembly.

Beyond the effect on the accuracy of phasing, the density of parental information provided for GT-Trio also affects the assembly contiguity and completeness. Haplotypes assembled with GT-Trio using lower density panels of array SNPs as input show significantly lower levels of fragmentation and fewer misassemblies between haplotypes than assemblies generated with higher-density parental information. The absence of haplotype-specific signals can simplify assembly through hard-to-assemble graph regions, as the assembly algorithm can choose the best path through the graph to preserve contiguity. However, it is important to not interpret the improved contiguity as genuine improvement of assembly quality, as it could rather reflect graph simplification and haplotype collapse disrupting correct haplotype phasing. Together these findings demonstrate how the density of parental information provided to GT-Trio influences both phasing accuracy and assembly contiguity. Understanding this balance is essential when considering the broader implementation of GT-Trio, as the optimal source and density of parental information will depend on the genomic resources available for a given population and the specific objectives of an assembly project.

The GT-Trio pipeline has proven its applicability for haplotype assembly in NR cattle and should be evaluated in other breeds and species where a high-quality breed-specific reference and imputation panel is available. Required reference and imputation resources may not be readily available in all species and populations, which may limit the applicability of the pipeline outside well-characterized livestock and aquacultural populations. However, our results suggest that GT-Trio remains effective with limited resources. Lower-density panels of array SNPs include enough haplotype-specific information to substantially improve haplotype phasing compared to trio-free assembly and can be used when high density panels of imputed sequence variants are not available. Similarly, using the more distant Hereford reference genome produced results only in a minor reduction of assembly quality and phasing accuracy, suggesting that while results are affected by reference quality and genetic similarity, the GT-Trio approach remains robust to moderate levels of reference divergence and can be used in population lacking population-specific reference. However, the sensitivity to increasing genomic divergence between the reference genome and the sample genomes has to be investigated before any general advice can be given regarding appropriate choice of reference.

GT-Trio can be used as a complementary approach to conventional trio-binning if parent individuals are unavailable for sequencing, enabling haplotype resolved assembly of a larger selection of genomes. Scalable strategies for haplotype-resolved assembly are especially valuable in the context of pangenome construction, where multiple assemblies are combined into a graph representing all the genetic diversity within a species or breed (37, 38). Haplotype-resolved assemblies are necessary to capture the full spectrum of structural variants and complex alleles (1, 39). A successful pangenome project has to start with biological design, rather than sequencing alone and should include samples representing as much diversity as possible. GT-Trio can facilitate the inclusion of a wider range of individuals with rare or particularly interesting phenotypes, which ultimately enables a more accurate and sensitive characterisation of structural variation, haplotype diversity, and functional alleles underlying phenotypes of interest.

## Declarations

### Ethics approval and consent to participate

Not applicable

### Consent for publication

Not applicable

### Availability of data and materials

The GT-Trio pipeline is publicly available at https://github.com/theahettasch/GT-Trio. Sequencing data from three Norwegian Red cattle trios used in this study is previously described by Hettasch et al., 2026, and is available on ENA under project ID PRJEB105120. Supplementary material is provided in Additional_file_1.docx, Additional_file_2.docx, Additional_file_3.docx and Additional_file_4.xlsx.

## Authors’ contributions (CRediT)

**TH:** Conceptualization, Methodology, Software, Validation, Formal analysis, Investigation, Data Curation, Writing – Original Draft, Visualisation. **AB:** Conceptualization, Methodology, Resources, Writing – Review & Editing. **MK:** Conceptualization, Methodology, Writing – Review & Editing. **HG:** Software, Methodology, Investigation, Data Curation, Writing – Review & Editing. **DV:** Conceptualization, Methodology, Writing – Review & Editing, Supervision, Project Administration, Funding acquisition.

## Acknowledgements

Not applicable

## Additional files

### Additional file 1, Methods S1

Format: .docx

Title: Parameters used to run GT-Trio.

Description: Parameters used to run the GT-Trio pipeline. They are defined in the configuration file (config.yaml).

### Additional file 1, Methods S2

Format: .docx

Title: Conventional trio-binning and trio-free assembly of NR haplotypes

Description: Code for running conventional trio-binning and trio-free assembly of NR haplotypes with Hifiasm.

### Additional file 1, Methods S3

Format: .docx

Title: Evaluation of accuracy of haplotype-separation with Meryl and Merqury

Description: Code used to evaluate accuracy of haplotype-separation based on k-mers with Meryl and Merqury.

### Additional file 1, Methods S4

Format: .docx

Title: Repeat annotation of assembly gaps and hapmer errors

Description: Code used to annotate the repeat content of genomic regions adjacent to assembly gaps and hapmer errors.

### Additional file 2, Figure S1

Format: .docx

Title: Centromere annotation of NR reference genome

Description: Annotation of centromere lengths in NR reference genome based on enrichment of bovine satellites.

### Additional file 2, Figure S2-S3

Format: .docx

Title: Size imbalance between haplotype assemblies

Description: Analysis of size imbalance between paternal and maternal haplotypes assembled with GT-Trio and the effect of switching the order of haplotype assignment for assembly.

### Additional file 2, Figure S4

Format: .docx

Title: Size and repeat content of unplaced contigs

Description: Analysis of size and repeat content of unplaced contigs constructed with conventional trio-binning and GT-Trio, and the effect of switching the order of haplotype assignment for assembly.

### Additional file 2, Figure S5

Format: .docx

Title: Misassembly between haplotypes in homozygous region

Description: Synteny plot between haplotypes (Bta14) showing larger regions of sequence, covering BUSCO genes, missing from one haplotype and present in the other. Figure also shows mapping of hapmers to misassembled region.

### Additional file 3, Table S1-S2

Format: .docx

Title: Informative SNPs across genotype panels

Description: Categorisation of SNP as informative, possible informative and not informative for haplotype-separation based on the combination of parent genotypes (Table S1). Number and proportion of informative, possibly informative and not informative SNPs across panels with sequence variants and array SNPs (Table S2).

### Additional file 4, Data S1

Format: .xlsx

Title: Offspring k-mers

Description: Number of paternal hapmers, maternal hapmers, shared k-mers and offspring-specific k-mers found in offspring-kmers across three NR trios.

### Additional file 4, Data S2

Format: .xlsx

Title: Assembly statistics

Description: Assembly statistics of NR haplotypes assembled with conventional trio-binning, trio-free assembly and GT-Trio.

### Additional file 4, Data S3

Format: .xlsx

Title: Assembly statistics t-test

Description: Paired t-tests comparing different assembly statistics of haplotypes assembled with conventional trio-binning and GT-Trio.

