## Additional_file_1 for "Haplotype assembly without parental sequencing: Genotype-based trio-binning (GT-Trio)"

**Supplementary Methods S1**

Parameters used to run GT-Trio.

The following parameter values were used to run the GT-Trio pipeline. For read simulation (step 2) a *read_length* of 150 bp and a *read_depth* of 30 was chosen to mimic the parental Illumina short-reads used for conventional trio-binning. *No error_rate, mutation_rate, indel_fraction* or *indel_extension* was applied to avoid introducing any errors to the simulated reads. For k-mer dictionary construction, a k-mer size of 21 was chosen as recommended by Koren et al. (2018). The minimum confidence scores *grouping_conf, location_conf and orientation_conf* were set to 0.7, 0.6 and 0.7 respectively to ensure confident scaffolding of contigs.

```
### Step 2: Read simulation parameters
read_length: "150"
read_depth: "30"
genome_size: "2800000000"
error_rate: "0"
mutation_rate: "0"
indel_fraction: "0"
indel_extension: "0"

### Step 3: K-mer dictionary construction parameters
yak_kmer: "21"
yak_bloom_bits: "37"

### Step 5: Scaffolding parameters
grouping_conf: "0.7"
location_conf: "0.6"
orientation_conf: "0.7"
```

**Supplementary Methods S2:**

Conventional trio-binning and trio-free assembly of NR haplotypes

Haplotype assembly of NR offspring genomes with conventional trio-binning was performed with Hifiasm (v0.24.0) (1). K-mer dictionaries were created from parental Illumina short-reads with Yak (2) using a bloom filter size (-b) of 2^37^, 4 threads (-t) and a k-mer size (-k) of 21.

yak count -o <paternal_count.yak> -b37 -t4 -k21 <paternal_reads.fq.gz>

yak count -o <maternal_count.yak> -b37 -t4 -k21 <maternal_reads.fq.gz>

Haplotype-resolved assemblies were generated from offspring ONT reads using Hifiasm in ONT mode (--ont), and parental k-mer dictionaries provided as input:

hifiasm --ont -o <offspring.asm> -1 <paternal_count.yak> -2 <maternal_count.yak> <offspring_reads.fq.gz>

Offspring ONT reads were also assembled without parental input. In trio-free assembly mode, Hifiasm produces two partially phased assemblies (offspring.asm.bp.hap1.p_ctg.gfa, offspring.asm.bp.hap2.p_ctg.gfa) based on local overlap between offspring reads.

hifiasm --ont -o <offspring.asm> <offspring_reads.fq.gz>

**Supplementary Methods S3**

Evaluation of accuracy of haplotype-separation with Meryl and Merqury

21-mer dictionaries were built from offspring ONT reads and parental Illumina short-reads with *Meryl* (v1.4.1) (3):

meryl k=21 count <paternal_illumina_reads.fq.gz> output <paternal.meryl>

meryl k=21 count <maternal_illumina_reads.fq.gz> output <maternal.meryl>

meryl k=21 count <offspring_reads.fq.gz> output <offspring.meryl>

Haplotype-specific k-mers (hapmers) were identified as the subset of offspring k-mers found uniquely in either the paternal of maternal set of k-mers with *Merqury* (v1.3) (3):

sh $MERQURY/trio/hapmers.sh <paternal.meryl> <maternal.meryl> <offspring.meryl>

Paternal and maternal hapmers were mapped to the paternal and maternal haplotype assemblies to assess the accuracy of haplotype separation. The number of paternal and maternal hapmers found on each chromosome was counted and visualised in a blob plot:

sh $MERQURY/trio/hap_blob.sh <paternal.meryl> <maternal.meryl> <paternal_assembly.fasta> <maternal_assembly.fasta> <out_prefix>

The hap_blob.sh script outputs a file called *out_prefix.hapmer.count* which includes the number of paternal and maternal hapmers observed in each chromosome across the paternal and maternal haplotype assemblies. These counts were used to calculate the *Hamming error rate:*

$$Hamming error rate= \frac{misassigned hapmers}{total hapmers-1}*100$$

**Supplementary Methods S4**

Repeat annotation of assembly gaps and hapmer errors

**Repeat annotation of NR reference genome**

A species-specific repeat database was built from the NR reference genome (GCA_963921495.1) with *RepeatModeler* (v2.0.7) (4), and subsequently used to perform a genome-wide repeat annotation with *RepeatMasker* (v4.1.7) (5):

BuildDatabase -name NR_DB NR_reference.fasta

RepeatModeler -database NR_DB -threads 10

RepeatMasker -pa 10 -gff -lib NR_DB-families.fa -dir NR_masked NR_reference.fasta

**Annotation of segmental duplications**

The annotation GFF output from *RepeatMasker* was converted to BED format and used to soft-mask the NR reference with *BEDtools* (v2.30.0) (6), which was then provided as input to *Biser* (v1.4) (7) for segmental duplication detection. Threads (-t) was set to 15 and Garbage Collection (GC) heap size (--gc-heap) was set to 32G to avoid extensive memory usage:

```
grep -v "^#" <NR_reference.fasta.out.gff> \
  | awk 'BEGIN{OFS="\t"}{print $1, $4-1, $5}' \
  > <NR_reference.repeats.bed>

bedtools maskfasta -soft \
  -fi <NR_reference.fasta> \
  -bed <NR_reference.repeats.bed> \
  -fo <NR_reference_softmasked.fasta>

biser -o <NR_SD> -t 15 --gc-heap 32G <NR_reference_softmasked.fasta>
```

**Liftover of contig breaks**

Contig breaks were inferred from the *RagTag* scaffolding output (ragtag.scaffold.agp and ragtag.scaffold.asm.paf) in three steps:

*Step 1 - Extract placed contig names and chromosomes:* Contigs placed on reference chromosomes were identified from the AGP file. Contig names (starting with h1tg or h2tg) and the NR reference chromosome (prefix Bta) they are placed on were retrieved from the AGP file. The *_RagTag* suffix was removed from the chromosome names:

awk -F'\t' 'BEGIN { OFS = "\t" }

$1 ~ /^Bta/ && $6 ~ /h[12]tg/ {

gsub("_RagTag", "", $1);

print $1, $6

}' ragtag.scaffold.agp > placed_contigs.txt

*Step 2 - Retrieve contig mapping positions:* For each placed contig, the corresponding alignment records were extracted from the PAF file. Only alignments where the contigs map to their assigned reference chromosome were retained:

awk -F'\t' 'BEGIN { OFS = "\t" }

NR==FNR {

contigs[$2]=$1; next

}

($1 in contigs && contigs[$1] == $6) {

print contigs[$1], $1, $6, $8, $9

}' placed_contigs.txt ragtag.scaffold.asm.paf > placed_contig_maps.txt

*Step 3 – Extract contig breaks:* For each contig, the minimum and maximum mapping positions across all alignment records were extracted to define the contig breaks. The output file contig_breaks.bed includes the reference chromosome, the placed contig name and the minimum and maximum mapping position.

```
awk -F'\t' 'BEGIN { OFS = "\t" }
{
	key = $1 FS $2;
    	min_pos = ($4 < $5) ? $4 : $5;
    	max_pos = ($4 > $5) ? $4 : $5;

    	if (!(key in min_map) || min_pos < min_map[key]) min_map[key] = min_pos;
    	if (!(key in max_map) || max_pos > max_map[key]) max_map[key] = max_pos;
}
END {
    for (k in min_map) {
        split(k, parts, FS);
        print parts[1], min_map[k], max_map[k], parts[2];
    }
}
' placed_contig_maps.txt | sort -k1,1 -k2,2n > contig_breaks.bed
```

**Liftover of hapmer error positions**

Hapmer error positions were lifted over from individual haplotype assemblies to the NR reference genome in three steps:

*Step 1 – Get BED file with hapmer error sites:* Hapmer error positions in in individual haplotype assemblies were identified using the *phase_block.sh* script from *Merqury* which outputs a file called *out_prefix.sort.bed* including all hapmer sites, both correct and erroneous. Erroneous hapmers positions are extracted from this file with awk:

sh $MERQURY/trio/phase_block.sh <assembly.fasta> <paternal.meryl> <maternal.meryl> <out_prefix>

awk -v hapmer_err="<roneous_hapmer_name>" '$6 == hapmer_err' out_prefix.sort.bed > hapmer_errors.bed

*Step 2 – Whole-genome alignment and CHAIN file conversion:* Each haplotype assembly was aligned to the NR reference genome using *minimap2 (8)*. Only same-chromosome alignments were retained. The PAF file output from was further converted to a CHAIN file with awk:

minimap2 -x asm5 -t 8 --cs=long <NR_haplotype.fasta> <NR_reference.fasta> > <NR_haplotype.paf>

sort -k6,6 -k8,8n <NR_haplotype.paf> > <NR_haplotype.sorted.paf>

awk '$1 == $6' <NR_haplotype.sorted.paf> > <NR_haplotype.same.sorted.paf>

awk '

BEGIN {

chain_id = 1

}

$5 == "+" {

print "chain 0", $1, $2, $5, $3, $4, $6, $7, "+", $8, $9, chain_id

print ($4 - $3), 0, 0

print "" chain_id++

}

' <NR_haplotype.same.sorted.paf> > <NR_haplotype.chain.txt>

*Step 3 – Liftover of hapmer error positions:* Hapmer error positions identified by Mercury in step 1 were lifted over to NR reference coordinates using *CrossMap (9)*:

CrossMap bed <NR_haplotype.chain.txt> \

<hapmer_errors.sort.bed> \

<hapmer_errors_liftover.bed>

**Intersection of contig breaks and hapmer error positions with repeat annotation**

BED files with contig break and hapmer error positions (contig_end.bed, hapmer_error_liftover.bed) were intersected with the repeat annotation from *RepeatMasker* (NR_reference.fasta.tsv) and the annotaton of segmental duplications from *Biser* (NR_SD.txt) using the R script below. Positions within ± 100 bp of an annotated repeat were considered repeat-associated. Additionally, positions falling within the first 5 Mb of an autosome were classified as centromeric.

```
### Import libraries
library(dplyr)
library(tidyr)

### Function: Import bed
import_bed <- function(file){
  df <- read.csv(file, sep = "\t", header = FALSE)
  df <- subset(df, select = c(1, 2, 3))
  colnames(df) <- c("Chr", "Start", "End")
  df <- pivot_longer(df, cols = c(Start, End), names_to = "Type", values_to = "Position")
  return(df)
}

### Function: Check if position is in repeat
is_position_in_repeat <- function(chrom, pos, repeats, l) {
  any(
    repeats$Chr == chrom &
    pos >= repeats$Start - l &
    pos <= repeats$End + l
  )
}

### Function: Annotate repeats
annotate_repeats <- function(repeat_type, df, repeats_df, l) {
  repeat_type_df <- repeats_df[grepl(repeat_type, repeats_df$Repeat_type), ]
  df[[repeat_type]] <- mapply(function(chrom, pos) {
    is_position_in_repeat(chrom, pos, repeat_type_df, l)
  }, df$Chr, df$Position)

  return(df)
}

### Function: Annotate repeats and centromeres
annotate_all <- function(types, df, repeats_df, l){

  for (t in types){
    df <- annotate(t, df, repeats_df, l)
  }
  return(df)
}

### Function: Count repeat types
count_repeat_types <- function(df){
  #Define repeat categories
  repeat_cat <- c("Centromere", "Low_complexity", "Simple_repeat", "LINE", "SINE", "LTR", "SD", "Unknown")

  # Add No_repeat column
  df$No_repeat <- !apply(df[repeat_cat], 1, any, na.rm = TRUE)

  # Update repeat_cols to include No_repeat
  repeat_cat <- c(repeat_cat, "No_repeat")

  # Count TRUEs in each column
  repeat_counts <- colSums(df[repeat_cat], na.rm = TRUE)

  # Convert to dataframe for plotting
  repeat_counts_df <- data.frame(
  repeat_type = names(repeat_counts),
  count = as.numeric(repeat_counts))

  return(repeat_counts_df)
}

### Import dataset with repeat annotations from RepeatMasker
repeats_df <- read.csv("NR_reference.fasta.tsv", sep = "\t", header = FALSE)
repeats_df <- repeats_df[grepl("Bta", repeats_df[[5]]), ]
repeats_df <- subset(repeats_df, select = c(V5, V6, V7, V11))
colnames(repeats_df) <- c("Chr", "Start", "End", "Repeat_type")
repeats_df$Start <- as.numeric(repeats_df$Start)
repeats_df$End <- as.numeric(repeats_df$End)

### Import dataset with SD annotations from Biser
SD_df <- read.csv("NR_SD.txt", sep = "\t", header = FALSE)
SD_df_1 <- subset(SD_df, select = c(V1, V2, V3))
colnames(SD_df_1) <- c("Chr", "Start", "End")
SD_df_2 <- subset(SD_df, select = c(V4, V5, V6))
colnames(SD_df_2) <- c("Chr", "Start", "End")
SD_df <- rbind(SD_df_1, SD_df_2)
SD_df$Repeat_type <- "SD"

### Import centromere positions
centromeres_df <- read.csv(“centromere_length.txt”, sep = ”\t”, header = TRUE)
centromeres_df$Repeat_type <- “Centromere”

### Combine all repeat annotations
ALL_repeats_df <- rbind(repeats_df, SD_df)

# Annotate Repeats and SDs
limit <-100 #adjecancy to repeat (bp)
repeat_types <- c("Low_complexity", "Simple_repeat", "LINE", "SINE", "LTR", "Unknown", "SD")

asm_bed <- import_bed("contig_breaks.bed/hapmer_errors_liftover.bed")
asm_df <- annotate_all(repeat_types, asm_bed, ALL_repeats_df, limit, centromere_length)

# Count repeat types
asm_counts <- count_repeat_types(asm_df)
```
