## Additional_file_2 for "Haplotype assembly without parental sequencing: Genotype-based trio-binning (GT-Trio)"

**Supplementary Figure S1**

Centromere annotation of NR reference genome

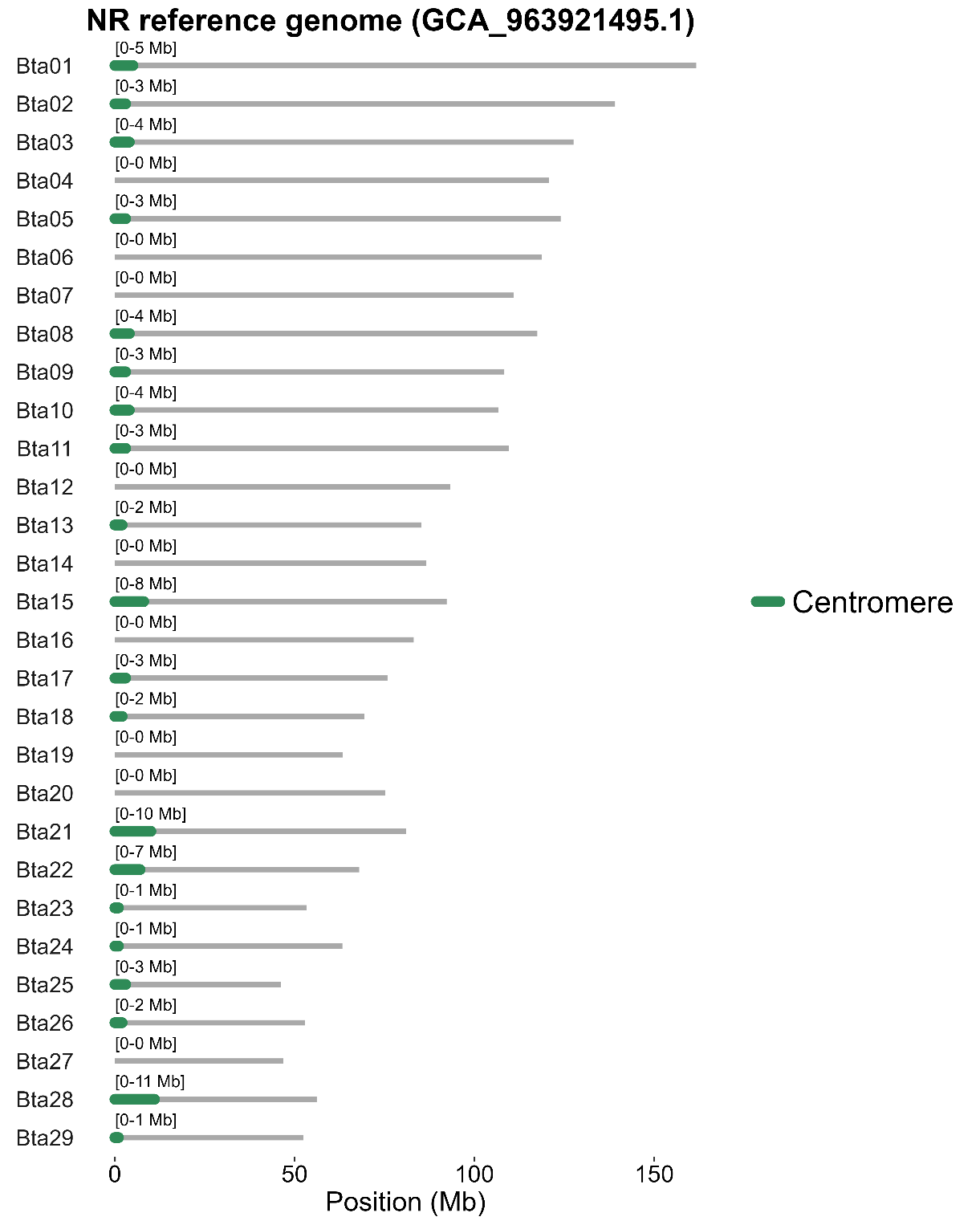

**Figure S1:** Centromere annotation of all acrocentric autosomes in the NR reference genome. Centromeres are regions significantly enriched in bovine satellites compared to the genome average. Centromeres are identified in 20 of the 29 NR autosomes and have a length of 1-11 Mb.

**Supplementary Figure S2-S3**

Size imbalance between haplotype assemblies

In the GT-Trio pipeline, the paternal data is provided as input for the assembly of haplotype 1 and the maternal data is provided as input for the assembly of haplotype 2. To assess if the order of parental assignment affects assembly size, assembly of NR haplotypes was repeated with maternal provided for assembly of haplotype 1 and paternal data for assembly of haplotype 2:

hifiasm --ont -o offspring.asm -1 maternal.yak -2 paternal.yak offspring_reads.fq.gz

In assemblies constructed with GT-Trio using array SNPs as input, the maternal contig-level assembly size is consistently larger than the paternal when the maternal data is assigned to haplotype 1 (Figure S2). This suggests that the assembly size imbalance is primarily driven by the order of haplotype assignment, rather than any inherent biological differences between the paternal and maternal haplotypes. However, when sequence variants are used as input, the paternal assemblies remain consistently larger than the maternal, regardless of the order in which the parental data is provided (Figure. S2). All trio sires used in this study were part of the reference population of short-read sequenced individuals used for imputation of sequence variants. One possible explanation is that paternal genotypes are more accurately imputed in our dataset, which could in turn influence haplotype assignment and contig retention, although this was not directly evaluated here.

After scaffolding against the NR reference genome no significant difference in length is observed between the paternal and maternal haplotype assemblies (Figure S3). The surplus of sequence present at the contig level ends up as unplaced contigs.

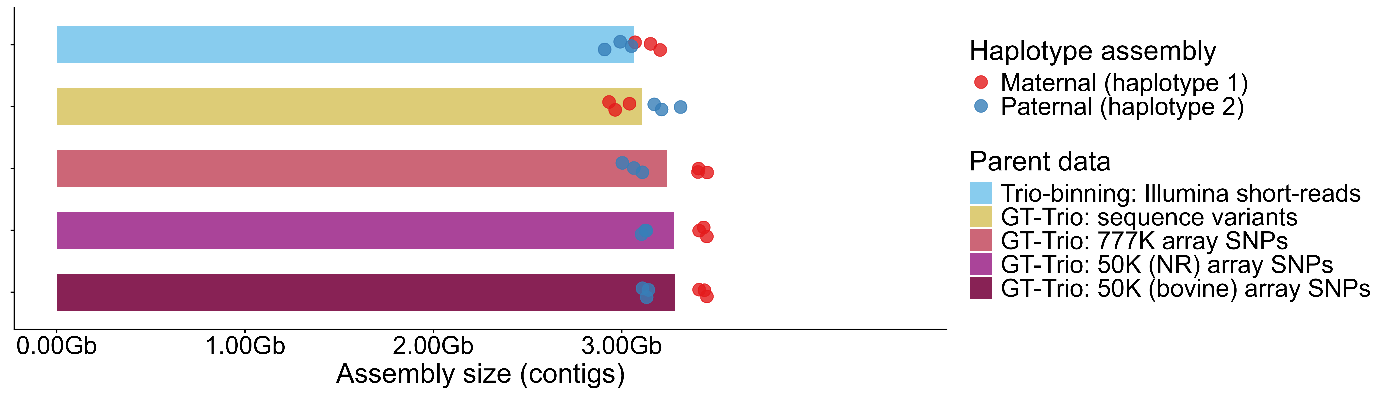

***Figure S2:*** *Contig-level assembly size of haplotypes assembled with maternal assembly considered haplotype 1 and paternal assembly considered haplotype 2.*

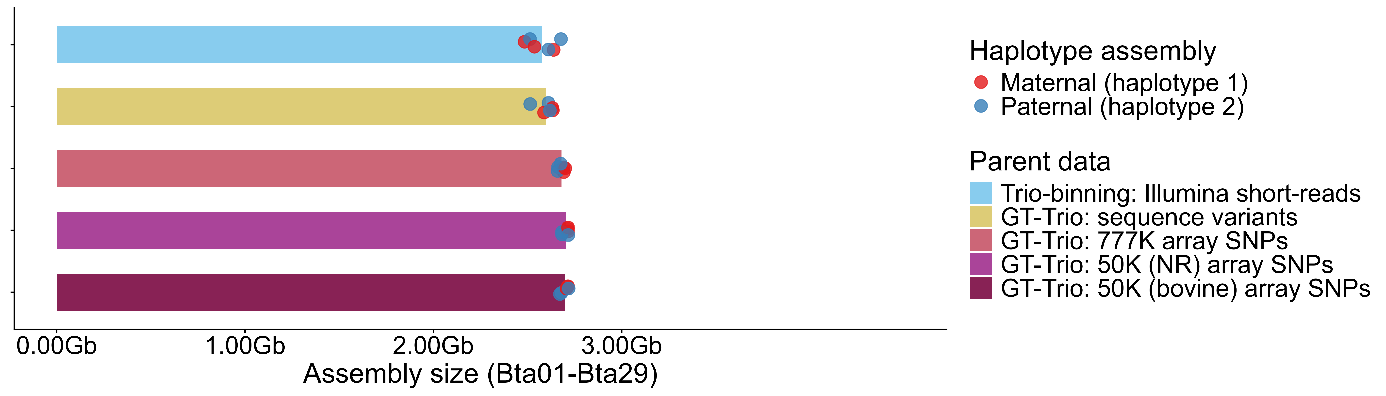

***Figure S3:*** *Chromosome-level assembly size of haplotypes assembled with maternal assembly considered haplotype 1 and paternal assembly considered haplotype 2.*

**Supplementary Figure S4**

Size and repeat content of unplaced contigs

To investigate the size and repeat content of unplaced contigs they were mapped to the NR reference genome with Minimap2 (1) and the number of repetitive seeds per unplaced contig was extracted from the output:

minimap2 -t 8 -x asm5 --secondary=no assembly.fasta unplaced_contigs.fasta > unplaced_contigs.sam

awk '{print $1, $2, $18}' unplaced_contigs.sam | sort -u > repetitive_seeds.txt

Figure S4 shows the length and repeat content of all unplaced contigs. Most unplaced contigs are short (<5 Mb) and highly repetitive (>50% harbouring repetitive seeds). Across assemblies constructed with GT-Trio using array SNPs as parental input, haplotype 1 includes a higher number of short and highly repetitive contigs than haplotype 2 (Figure S4), explaining why haplotype 1 is consistently larger than haplotype 2 at the contig-level. In assemblies constructed with GT-Trio using sequence variants as parental input there is also a higher number of short and highly repetitive contigs in haplotype 1 compared to haplotype 2. However, any long and highly repetitive contigs are only present in the paternal assembly, explaining why the paternal haplotypes are consistently larger than the maternal haplotype in these assemblies.

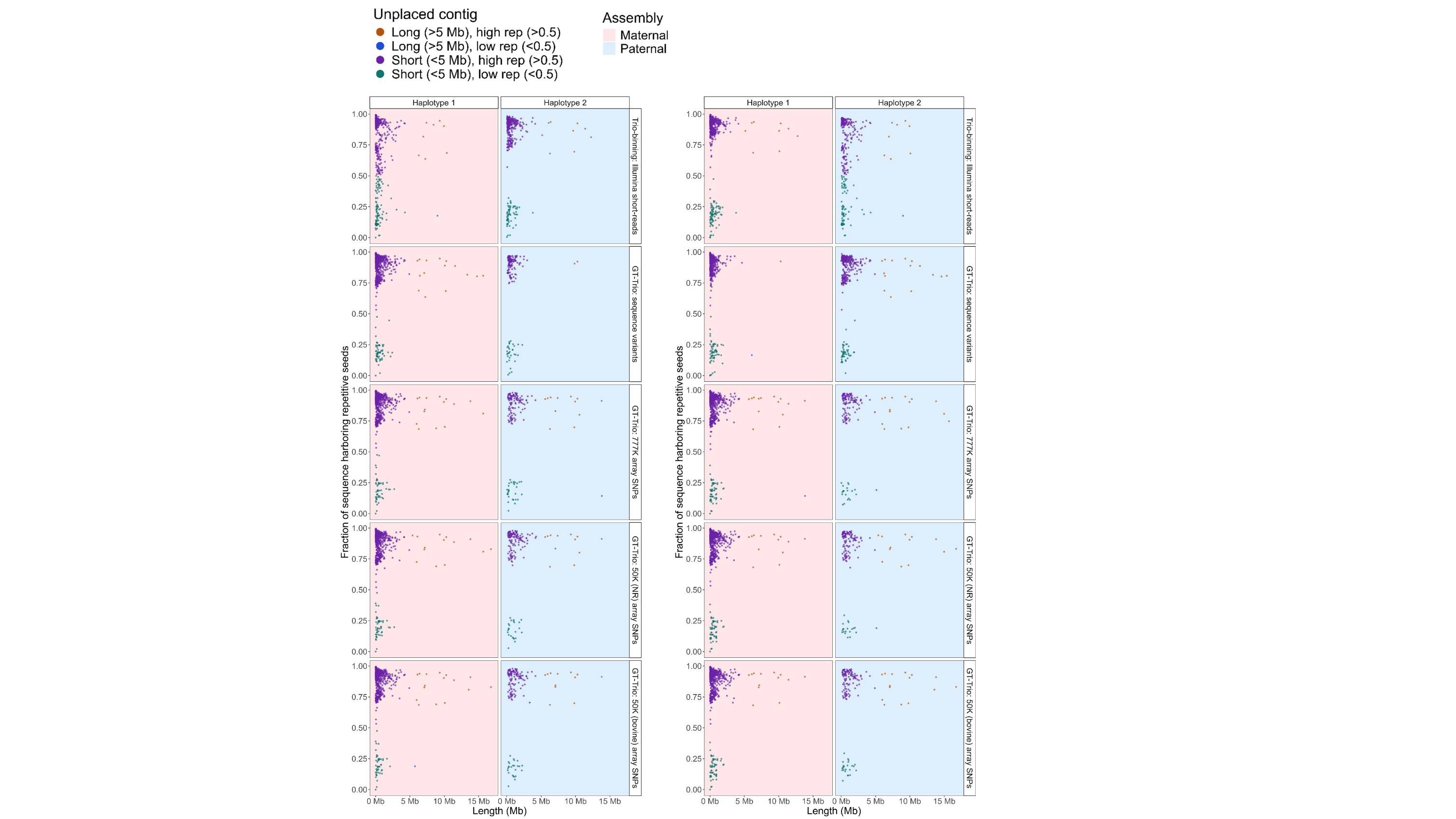

**Figure S4:** Length (Mb) and repeat content (Fraction of sequence harbouring repetitive seeds) of unplaced contigs extracted from assemblies constructed with conventional trio-binning and GT-Trio. In the left panel the paternal assembly is considered haplotype 1 and the maternal assembly is considered haplotype 2. In the right panel the order of haplotype assignment has been switched. Unplaced contigs are divided into four categories based on their length and repeat content: Long (>5 Mb) and high-rep (>0.5), long (>5 Mb) and low-rep (<0.5), short (<5 Mb) and high-rep (>0.5) and short (<5 Mb) and low-rep (<0.5).

**Supplementary Figure S5**

Misassembly between haplotypes in homozygous region

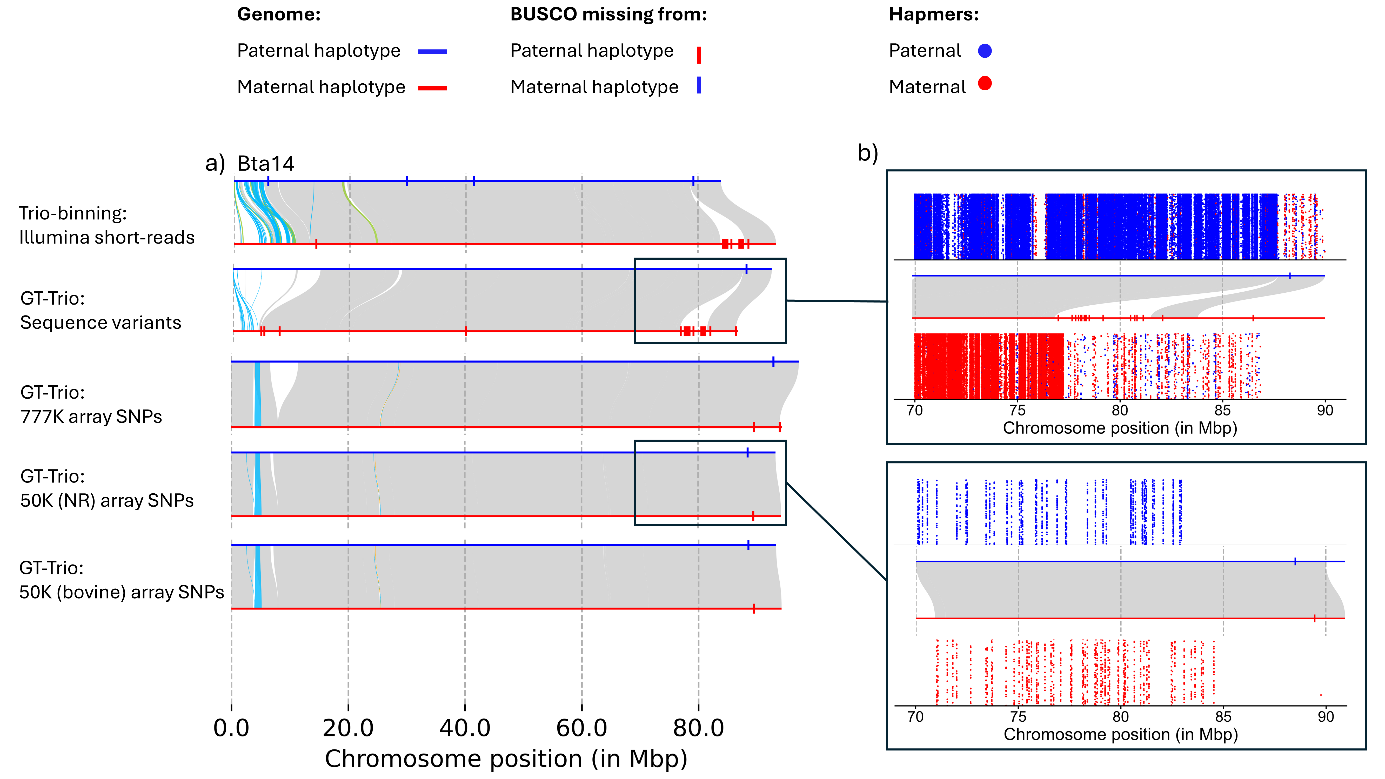

**Figure S5:** a) Synteny between haplotypes (Bta14) assembled with conventional trio-binning and GT-Trio for NR trio 1. In the paternal haplotype assembled with high-density parental information from short-reads or sequence variants a larger region of sequence covering several BUSCO genes is missing. The region is included in the maternal haplotype. The misassembly is not observed in haplotypes assembled with GT-Trio using lower-density panels of array SNPs. b) Mapping of hapmers recovered from sequence variants show low hapmer density and high hapmer ambiguity in misassembled region. Mapping of hapmers recovered from 50K NR-specific array SNPs show no hapmer ambiguity.
