## Additional_file_3 for "Haplotype assembly without parental sequencing: Genotype-based trio-binning (GT-Trio)"

**Supplementary Table S1-S2**

Informative SNPs across genotype panels

**Table S1:** Informativeness of parental SNP genotypes for separation of offspring haplotypes. Parental SNP genotypes can be informative, possibly informative or not informative for separation of offspring haplotypes depending on the combination of parent genotypes.

| Parent genotype | Offspring genotype | Informativeness for haplotype separation |
| --- | --- | --- |
| AA, BB | AB | Informative |
| AA, AB | AA/AB | Possibly informative* |
| AB, AB | AA/BB/AB | Possibly informative** |
| AA, AA | AA | Not informative |

*Informative if offspring is heterozygous

**Informative if two or more HET alleles with diverging phase in parents are covered by same k-mer.

**Table S2:** Number and proportion of parental SNPs with informative, possibly informative and non-informative combinations of genotypes across sequence variants and array-based SNP panels used in this study.

| SNP panel | Trio | Informative | Possibly informative | Not informative |
| --- | --- | --- | --- | --- |
| Sequence variants | 1 | 694 910 (3.3%) | 6 349 417 (30.3%) | 13 902 517 (66.4%) |
| Sequence variants | 2 | 791 078 (3.8%) | 6 287 489 (30.0%) | 13 868 277 (66.2%) |
| Sequence variants | 3 | 644 115 (3.1%) | 6 485 784 (31.0%) | 13 816 945 (66.0%) |
| 777K array SNPs | 1 | 44 573 (7.1%) | 339 741 (54.3%) | 241 766 (38.6%) |
| 777K array SNPs | 2 | 50 111 (8.0%) | 334 806 (53.5%) | 241 163 (38.5%) |
| 777K array SNPs | 3 | 42 207 (6.7%) | 344 689 (55.1%) | 239 184 (38.2%) |
| 50K (NR) array SNPs | 1 | 3 750 (7.7%) | 28 332 (58.2%) | 16 577 (34.1%) |
| 50K (NR) array SNPs | 2 | 4 324 (8.9%) | 28 210 (58.0%) | 16 125 (33.1%) |
| 50K (NR) array SNPs | 3 | 3 637 (7.5%) | 29 472 (60.6%) | 15 550 (32.0%) |
| 50K (bovine) array SNPs | 1 | 2 980 (6.6%) | 23 289 (52.0%) | 18 544 (41.4%) |
| 50K (bovine) array SNPs | 2 | 3 428 (7.6%) | 23 045 (51.4%) | 18 340 (40.9%) |
| 50K (bovine) array SNPs | 3 | 2 802 (6.3%) | 23 816 (53.1%) | 18 195 (40.6%) |
